# Energetics of substrate transport in proton-dependent oligopeptide transporters

**DOI:** 10.1101/2024.05.01.592129

**Authors:** Balaji Selvam, Nicole Chiang, Diwakar Shukla

## Abstract

The PepT_So_ transporter mediates the transport of peptides across biological membranes. Despite advancements in structural biology, including cryogenic electron microscopy structures resolving PepT_So_ in different states, the molecular basis of peptide recognition and transport by PepT_So_ is not fully elucidated. In this study, we employed molecular dynamics simulations, Markov State Models (MSMs), and Transition Path Theory (TPT) to investigate the transport mechanism of an alanine-alanine peptide (Ala-Ala) through the PepT_So_ transporter. Our simulations revealed conformational changes and key intermediate states involved in peptide translocation. We observed that the presence of the Ala-Ala peptide substrate lowers the free energy barriers associated with transition to the inward-facing state. Furthermore, we elucidated the proton transport model and analyzed the pharmacophore features of intermediate states, providing insights for rational drug design. These findings highlight the significance of substrate binding in modulating the conformational dynamics of PepT_So_ and identify critical residues that facilitate transport.

## Introduction

The cell membrane is a dynamic cellular structure that predominantly functions as a selective permeability barrier by blocking the free exchange of biological molecules between the cytoplasm and the external environment. Solute carrier transporters (SLC) are specialized carrier biomolecules involved in active exchange with ions, protons, amino acids, and nucleosides across the cellular membrane and serve as one of the most fundamental and highly regulated processes in living cells.^1,2^ SLC15 gene family members, peptide transporter 1, PepT_1_ (SLC15A1) and peptide transporter 2, PepT_2_ (SLC15A2) are expressed in intestine and kidney tissues and are actively involved in the uptake of dietary peptide molecules. These transporter proteins are targeted for efficient drug delivery strategies to improve the pharmacokinetic profiles of the drug molecules. ^3–6^ The peptide transporter belongs to proton-dependent oligopeptide transporter (POT) family, which are members of the major facilitator superfamily (MFS) that facilitate the transport of small groups of amino acids across the cell membrane. MFS family proteins function via an alternate access model that involves alternate accessibility to either side of the membrane via distinct inward-facing (IF), occluded (OC), and outward-facing (OF) conformational states.^7,8^

The modern drug delivery approach uses these transporter proteins, more specifically PepT_1_ and PepT_2_, due to wide expression in the small intestine and kidney, respectively, broad substrate specificity, and high transport capacity.^9–14^ For example, Valacyclovir, a PepT_1_-targeted prodrug, shows enhanced oral drug delivery and increases the bioavailability up to 50%.^15^ To optimize the design of drugs targeting POTs, it is crucial to understand the interdependence between conformational dynamics, substrate binding, and proton transport in POTs. While significant progress has been made, Li *et al*. explored general proton transport principles,^16^ Parker *et al*. focused on proton transport in POTs,^17^ and Lichtinger *et al*. investigated PepT_2_’s gating cycle with enhanced sampling molecular dynamics (MD). ^18^ However, a detailed understanding of how these factors work together in POTs remains to be fully elucidated. To this end, Markov state models (MSMs) are promising tools.^19^ By constructing a coarse-grained representation of a transporter’s conformational space, MD data analyzed using MSMs enable exploration of long timescales and identification of key transition states. This approach has been successfully applied to study the substrate translocation of various membrane transporters, including the serotonin transporter and the AtSWEET13 sugar transporter, revealing interactions between substrate binding and transporter gating.^20–23^ These successful applications in other transporters highlight the potential of MSMs for studying proton-substrate transport in POTs.

This study utilizes unbiased MD simulations and MSMs to explore the complete proton-substrate transport pathway of *Shewanella oneidensis* peptide transporter, PepT_So_. Specifically, we focus on how substrate binding influences the transporter’s structure and how these structural changes impact the pathway for proton transport. Despite the availability of crystal structures for human PepT_1_, human PepT_2_, horse PepT_1_, and rat PepT_2_, our study primarily utilizes the crystal structure of PepT_So_ as a general template to investigate the broader function of POTs.^24–28^ We identified the crucial residues which drive the structural transitions to various intermediate states, and estimated the free energy barriers between them. Our earlier study revealed the OF structure of the SCL15 gene family for the first time, and provided an opportunity to characterize how the substrate (di/tri-peptide) or prodrug (Valacyclovir) molecules are recognized and transported via peptide transporters. We also analyze the proton transport in context of the behavior of ionizable groups across three key protein conformations: the inward-facing (IF), occluded (OC), and outward-facing (OF) states. We show that protons involved in transport move through well-ordered water clusters. While our approach utilizes traditional MD, which does not explicitly model the proton transfer event itself, it allows us to gain valuable insights into the potential pathway for proton movement based on the behavior of these ionizable groups within the changing protein environment. Our findings elucidate how substrate and proton transport across the biological membrane, which may inform drug design targeting POTs.

## Methods

### MD simulation setup

All-atom molecular dynamics (MD) simulations were performed using Amber18.^29^ The OF structure PepT_So_ was used as a starting structure for MD simulation.^30^ PepT_So_ was embedded in a phospho lipid bilayer and solvated using TIP3P water molecules.^31^ 100mM concentration of Ala-Ala peptide (*∼* equivalent to 12 molecules) was added to the simulation box using Packmol. The MD system was neutralized using 150mM of salt concentration. MD system was built using the tleap program in AmberTools18.^29^ The default amberFF14SB^32^ was used for simualtion. MD systems were subjected to minimization for 20,000 steps using the conjugate gradient method. The MD systems were slowly heated from 0 to 300K over a period of 1ns in NVT ensembles. All production runs were conducted in NPT ensembles at 300K using a Langevin thermostat. The pressure was kept constant at 1atm using a Langevin barostat. The integration step size was set to 2 fs. The particle mesh Ewald method was used for long range electrostatics calculations and the SHAKE algorithm was used to constrain bonds containing hydrogen.^33^ The cutoff for long-range Van der Waals interactions was set to 10 Å. MD system was equilibrated for 50 ns and the productions runs were carried out over a period of *∼*85 *µ*s.

### Adaptive sampling

Proteins exist in a dynamic ensemble of conformations. However, observing the transition between these states can be challenging because they often occur on very long timescales, making them difficult to capture with short molecular dynamics simulations. To efficiently explore the conformational landscape of the protein, we employed an adaptive sampling approach. This method leverages reaction coordinates that capture the slowest processes on the protein’s free energy landscape and has been employed to obtain the complete free energy landscapes of membrane proteins including transporters.^3,30,34–37^ The adaptive sampling procedure involves three iterative steps: (1) running a set of short MD simulations from various initial protein structures, (2) clustering the protein conformations observed in these simulations using the K-means algorithm into distinct states according to structural metrics such as gating distances and substrate location in the translocation tunnel, and (3) seeding new MD simulations from strategically selected states based on a sampling criterion. In this study, we employed the least counts approach as our sampling criterion. Recently, several machine learning based adaptive sampling approaches have been reported.^38–40^ In this case, we already knew the key reaction coordinates involved in the transport cycle based on our previous simulation study of the PepT*_So_*.^30^ Therefore, the least count based adaptive sampling combined with the known reaction coordinates is expected to be provide similar efficiency as the current state-of-the-art adaptive sampling methods. This approach prioritizes conformations from clusters with lower population densities, aiming to overcome potential biases towards frequently observed states and ensure a more comprehensive exploration of the conformational space. It’s important to note that any bias introduced by the least counts approach is mitigated during the construction of the final Markov state model (MSM) by estimating the reverse transition probability matrix for transitions between all conformational states. In the limit of long simulation timescales, the sampling errors introduced by this approach are expected to be negligible. The reaction coordinates used in this study to guide the adaptive sampling process were specifically chosen to capture the key dynamic processes of interest. These coordinates included the distances between the extracellular and intracellular gating residues, which are crucial for regulating substrate access, and the z-coordinate of the alanine dipeptide (Ala-Ala) substrate molecule being transported. By focusing on these key metrics, we were able to efficiently explore the conformational changes associated with substrate translocation. Through this iterative process of exploration, clustering, and strategic seeding, we performed a total of 34 rounds of adaptive sampling simulations. The length of simulation per round is listed in Table S1.

### Markov state modeling

The simulation data was analyzed using MSMBuilder 3.6.^41^ To build an MSM, we construct a transition probability matrix between states, allowing us to determine the rate of transition from one state to other. 13 residue pair distances and z-coordinate of Ala-Ala peptide was used for the MSM construction. The number of clusters and appropriate lag time was obtained by using a hyperparameter tool, Osprey (Fig. S13). The lagtime of 20 ns was used for MSM construction (Fig. S13). Transition path theory (TPT) was used to obtain the most probable pathway for substrate translocation and residues that drive the conformational transition.^42,43^ The synthetic trajectories are obtained using kinetic Monte Carlo simulation on the MSMs, allowing us to determine the substrate driven conformational changes in PepT_So_.^44^

### Trajectory analysis

The post processing of the simulation data was performed using a python packages such as MDTraj,^45^ Pytraj^46^ and Cpptraj.^47,48^ HOLE program was used to plot the translocation pore radius plots. ^49^ The structure based pharmacophores are obtained using Cavpharmer. ^50^ The trajectories were analyzed and visualized using VMD1.9.4^51^ and pymol v 2.0.^52^

## Results and discussion

### Substrate-Induced Conformational Changes Drive Efficient Peptide Transport in PepT_So_

We utilized molecular dynamics (MD) simulation with a Markov state model (MSM) framework to study the effect of substrate transport on the conformational dynamics of PepT_So_. Building upon previous studies, the initial conformation for our simulations was chosen as the MD predicted OF state of PepT_So_.^30,53^ To investigate the transport mechanism of the Ala-Ala peptide, a concentration of 100mM (approximately 12 molecules) was added to the simulated system. Consistent with established experimental data, Asp316 in helix 7 was maintained in its protonated state^54^ and roughly 85 µs of simulation data was obtained using an adaptive sampling approach. ^55^

Our MSM analysis successfully elucidated the transport mechanism of the Ala-Ala peptide in PepT_So_. This analysis also identified the functionally relevant intermediate states involved in the translocation process. These states include the inward-facing (IF), occluded (OC), and outward-facing (OF) conformations. To investigate the energetics of this process, we generated a conformational free energy landscape plot (Fig.1). This plot visualizes the free energy barriers between the three key intermediate states. To construct this landscape, the MD data was projected onto two key distances: (1) the extracellular distance between Arg32 (TM1) and Asp316 (TM7) and (2) the intracellular distance between Ser131 (TM4) and Tyr431 (TM10). The MSM equilibrium probability distribution was used to weight the data points within the landscape to ensure that the plot accurately reflects the free energy of the simulated system.

**Figure 1:**
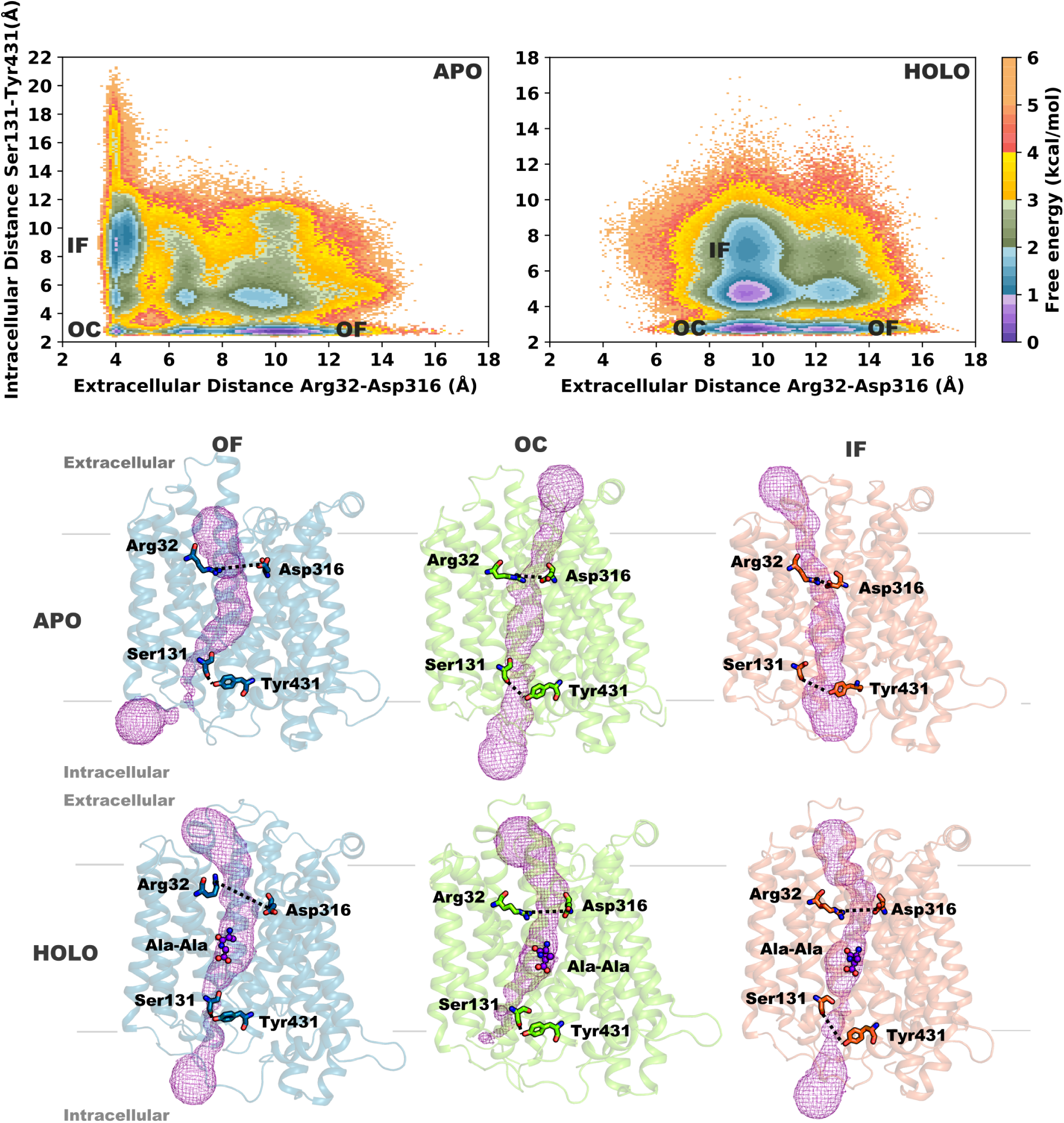
Free energy landscape for PepT_So_ in both apo (without substrate) and holo (with Ala-Ala peptide) states. The transitions between the inward-facing (IF), occluded (OC), and outward-facing (OF) states are depicted. Representative structures of IF, OC, and OF states for both apo and holo forms are shown. The extracellular gating residues (Arg32-Asp316) and intracellular gating residues (Ser131-Tyr431) are shown in sticks.

Our analysis of the free energy landscape and representative structures in Figure 1 provides valuable insights into the transport mechanism of the Ala-Ala peptide within the holo state (with substrate). The initial step involves substrate recognition by Asn22, Glu336, and Arg32 at the periplasmic side, stabilizing the OF state. This initial binding likely occurs with a relatively low free energy barrier (¡ 0.5 kcal/mol). As the Ala-Ala peptide moves towards the center of the translocation pore, PepTSo transitions to the OC state, characterized by a lower free energy compared to the OF state (¡ 0.5 kcal/mol). This transition coincides with a decrease in the distance between the extracellular gating residues, Arg32 (TM1) and Asp316 (TM7), to around 5-7 Å. This closure likely involves the movement of the extracellular domains in transmembrane helices TM1-2 and TM7-8. Further downward movement of the substrate triggers the opening of the intracellular cavity, allowing the peptide to exit the transporter. This final step is accompanied by the movement of the intracellular gating residue outwards, leading to the inward-facing (IF) state with a free energy of approximately 1.5 kcal/mol and an increased distance between the gating residues (10-11 Å).

Figure 1 also depicts the free energy landscape and representative structures for the apo state (without substrate) from our previous study.^30^ While the overall transport pathway remains similar, the presence of the substrate reduces the free energy barrier between the OC and IF states by roughly 1 kcal/mol. This suggests that substrate binding facilitates the release of the peptide from the transporter’s binding pocket. Furthermore, Figure 1 depicts representative structures of the IF, OC, and OF states for both apo and holo forms. Notably, the structures reveal a more open OF state in the holo form compared to the apo form. This observation is corroborated by the wider extracellular distance in the holo form, which is roughly 3 Å larger as shown in Figures 1 and S1. These findings together indicate that the presence of the Ala-Ala peptide within the transport tunnel widens the extracellular gate, likely promoting transition to the IF state and substrate release.

### Free Energy Landscape projected on tICs Reveals Key Slow Kinetic States

To gain deeper insights into the slow processes governing the transport mechanism, we employed time-independent component (tIC) analysis.^56^ This method projects the highdimensional MD data onto a lower-dimensional space spanned by the slowest collective motions of the system. The resulting free energy landscape, projected onto the first two tICs, is depicted in Figure 2. This landscape reveals three distinct minima, corresponding to kinetically slow states involved in the transport cycle. Figure 2 and S14 showcases representative structures and features, respectively, corresponding to each minimum in the free energy landscape. Interestingly, the first minimum represents the most stable state based on the free energy landscape. In this OF-OC state, the Ala-Ala peptide might be present within the translocation tunnel and potentially interacting with initial binding sites, but not yet fully secured at the specific orthosteric site within PepT_So_. This observation suggests a favorable initial recognition between the peptide and the transporter, followed by a low free energy barrier transition towards the second minimum, which represents the OC state. The second minima (OC) corresponds to a state where the Ala-Ala peptide is firmly bound at the orthosteric site within the occluded conformation of PepT_So_. This state suggests a crucial step where the translocation process pauses for efficient and specific substrate interaction before proceeding, as shown by the high free energy barrier from the second to the third minima, which depicts the inward-facing (IF) state. In this state, PepT_So_ adopts a conformation with the intracellular gate open, allowing the peptide to reach the intracellular end of the translocation tunnel. These findings from the tIC analysis complement the free energy landscape analysis from Figure 1 by providing a more detailed view of the slowest steps during the transport process. The low free energy barrier between the initial recognition (first minimum) and bound state (second minimum) suggests a favorable and efficient transition between these crucial steps. The identification of the OC state as a stable minimum highlights the importance of this state for proper substrate interaction before translocation proceeds.

**Figure 2:**
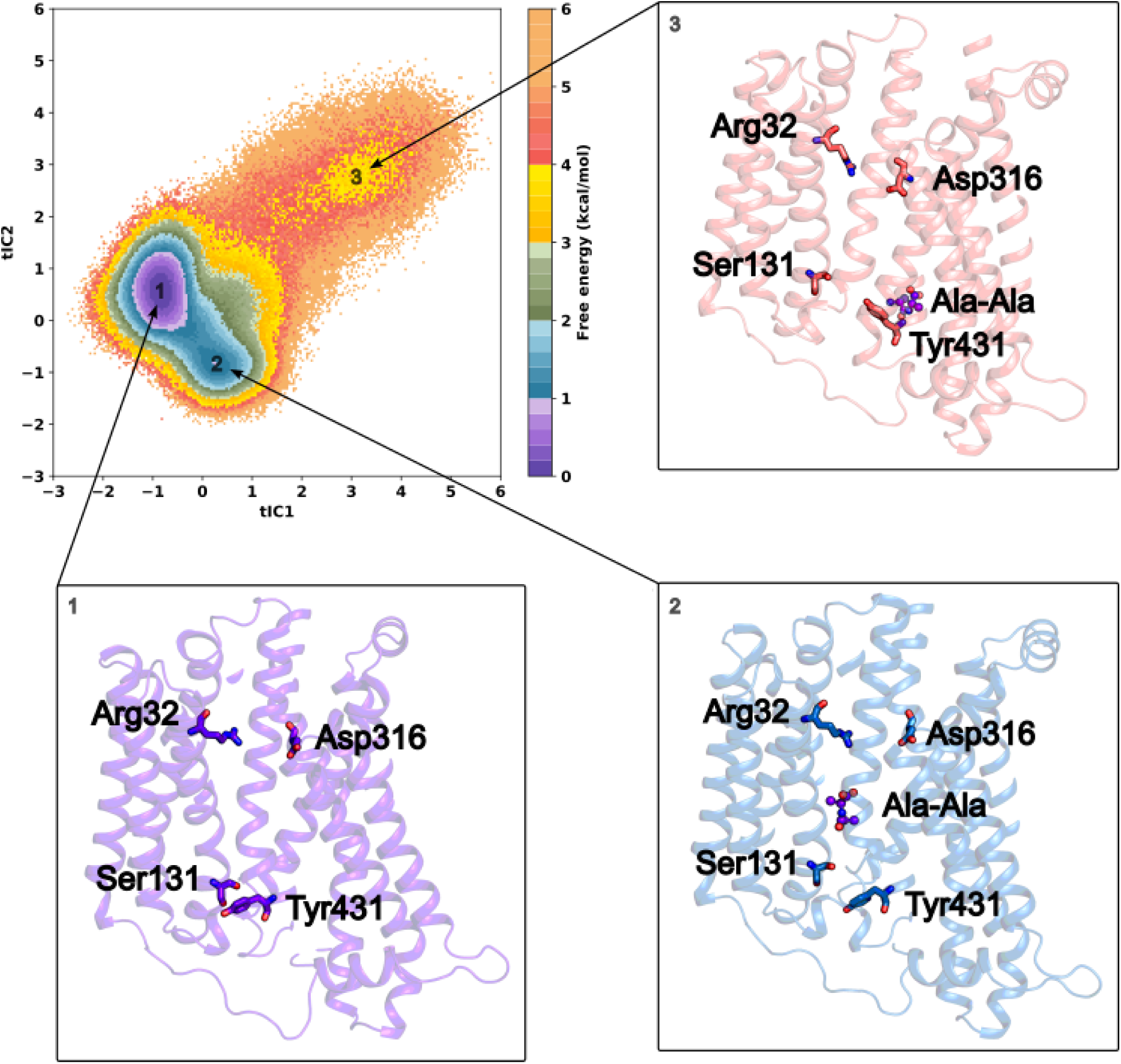
Free energy landscape of PepT_So_ projected onto the first two time-independent components (tICs), representing the slowest processes in the system. The transitions between the three kinetically slow states (OF-OC, OC, and IF) are depicted. The representative structures corresponding to the three minima in the free energy landscape are depicted. The first minimum (OF-OC) represents a conformation where the peptide may be present within the translocation tunnel but not bound to the orthosteric site. The second minimum (OC) represents PepT_So_ with the Ala-Ala peptide bound at the orthosteric site in an occluded conformation. The third minimum (IF) represents the inward-facing conformation of PepT_So_ with the Ala-Ala peptide positioned at the intracellular end of the translocation tunnel, signifying its release from the transporter.The extracellular gating residues (Arg32-Asp316) and intracellular gating residues (Ser131-Tyr431) are shown in sticks.

### TPT Identifies Key Interactions for Conformational Shifts in Peptide Transport

Using Transition Path Theory (TPT), we determined the likely pathway of the polypeptide substrate’s translocation mechanism through different functional states. TPT identifies the top flux pathways from the transition probability matrix to determine the critical residues involved in the transport pathway. The peptide molecule was recognized by Asn22, Arg32, Glu336 and Gln341 at the extracellular surface and enters the transporter (Fig. 3A, 3B). Arg32 coordinates with the negatively charged carboxylate group of Ala-Ala and initiates the transport from the extracellular surface to the center of the transporter. The mutation of Arg33 to alanine results in the disruption of proton-driven substrate transport in PepT_St_;^57^ in contrast, such an effect was not observed in PepT_So_.^17^ The primary amine of the peptide diffuses into the pore cavity and interacts with Gln341 and thereby facilitates transport (Fig. 3B). The polypeptide molecule moves down and forms additional polar contacts with Asp316, His61 and Tyr68 to stabilize the OF state (Fig. 3C). Site-directed mutagenesis experiments show that mutation of proton-binding residue Asp316 to alanine results in loss of function.^17,54,57,58^ His61 is conserved in human PepT_1_ and PepT_2_, hence playing a crucial role in transport.^17^ The peptide interaction involving Arg32 and Asp316 partially closes the extracellular surface as the distance between the gating residues decreases from *∼*13 Å to *∼*10 Å (Fig. 3D, 3E). The peptide molecule slowly diffuses to the center of the transporter and Arg25 forms an ion-pair interaction with peptide while Ser423 recognizes the amino terminal of the substrate (Fig. 3F). The diffusion of peptide inside the translocation pore was favored by conserved Tyr29 and Tyr68 residues (Fig. 3F, S2). Tyr29 and Tyr68 has been shown to play a vital role in determining the substrate specificity between the di-/tripeptide molecules.^57–59^ Arg25 at the central cavity belongs to the ExxERxxxY motif which is conserved across the mammalian POT family and mutation of this residue disrupts proton-driven peptide transport.^54,57^ At this juncture, the extracellular part of N- and C-terminal domains comes close to each other and decreases the gating residue distance to *∼*6-7 Å to obtain OC state (Fig. 3F). The predicted OC state shows good agreement with PepT_St_ structure (Fig. S3). The OC state is least stable compared to previous studies, as the protonated Asp316 decreases the strength of the hydrogen network with gating residues[10]. Further downward movement of the peptide substrate leads to opening of the intracellular cavity as the helices TM4-TM5 and TM10-TM11 move away from each other. Ser131 and Glu428 are involved in hydrogen bond interaction and stabilizes the peptide molecule in the translocation pore (Fig. 3G, 3H). Glu428 is conserved in human intestinal peptide transporters and is hypothesized to play crucial role in substrate driven proton transport. Finally, the peptide molecule interacts with Tyr431 and leaves the transporter (Fig. 3I). The intracellular side of the transporter opens to the cell as the distance between the gating residue increases up to *∼*10-12 Å to obtain the IF state (Fig. 3J).

**Figure 3:**
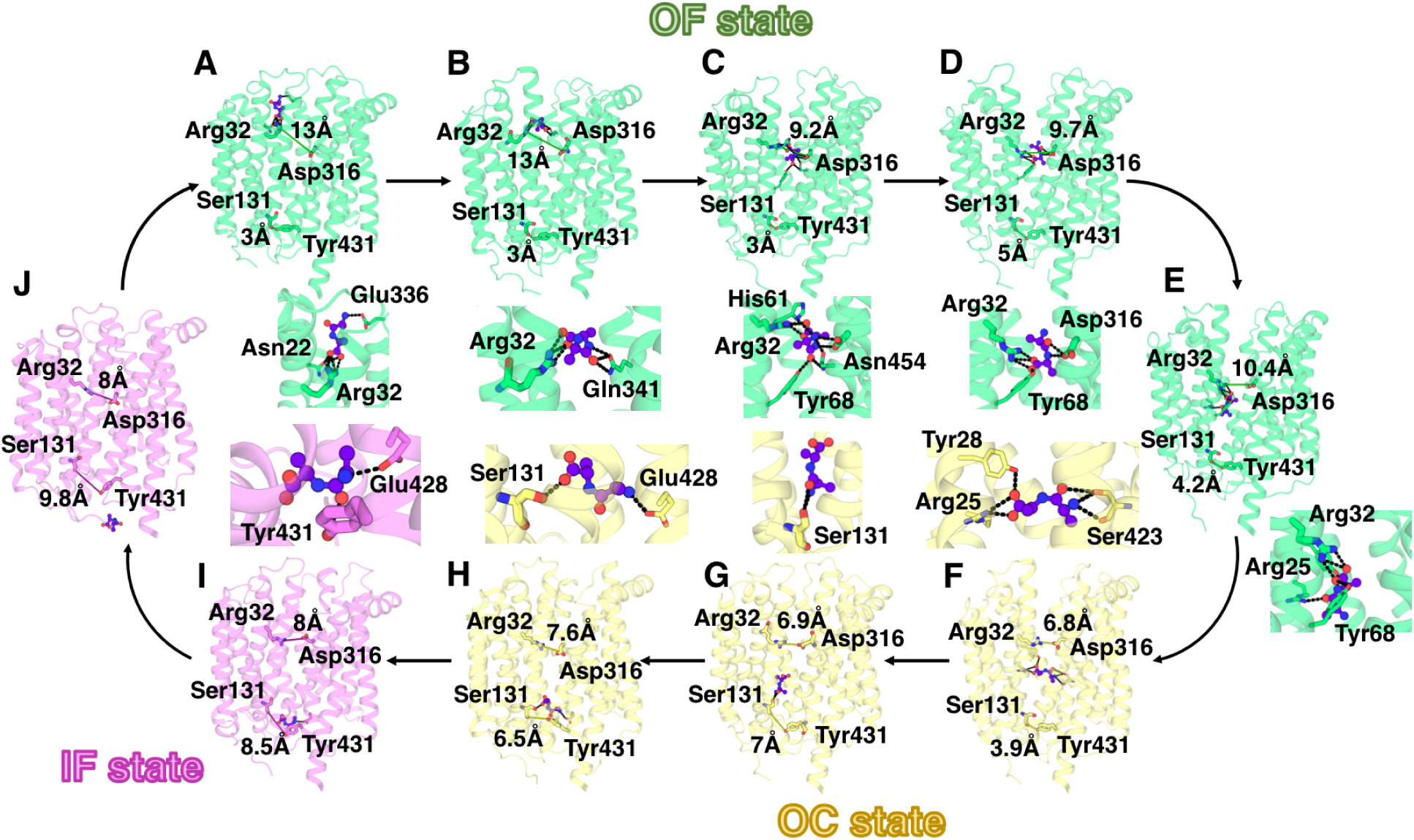
Substrate transport in PepT_So_. The OF, OC and IF states are represented as cartoons in green, yellow and magenta, respectively. Ala-Ala peptide was shown as ball and stick model and in purple color. The residues involved in interactions are shown as sticks for OF, OC and IF states

### Well-Organized Water Clusters Aid Proton Transport

Previous studies hypothesized the proton transport model in different peptide transporters.^17,54^ Three glutamates residues at the extracellular half are involved in proton transfer in PepT_St_.^17^ Parker *et al*., investigated the proton transfer using quantum mechanical studies and predicted Asp322 and Glu425 are exhibited important role in H+ transport.^17^ We analyzed the intermediate states such as OF, OC and IF in the simulation data and calculated the pKa values of ionizable groups using PROPKA to identify the possible proton transfer pathway.^60^ In OF state, Asp316 was predicted in protonated form while Glu419 and Glu428 exhibited deprotonated form. Glu419 is buried in the structure and forms extensive polar contact with Lys318 and Asn344 (Fig. S4). The equivalent residue Glu391 in E.coli peptide transporter plays a crucial role in active transport and mutation to glutamine decreases the substrate uptake. ^59^ At the cytoplasmic side, the negative charge on Glu428 is neutralized by a neighboring residue of opposite charge, Lys375 (Fig. S4). However, Asp316 is predicted as deprotonated in OC state and two glutamates has similar profiles as OF state (Fig. S5). In contrast, IF state shows major structural changes at the extracellular and intracellular side. Asp316 and Glu419 are aligned favorably for proton transfer (Fig.4, S6). Also, helix 10 undergoes major conformational change and rotates 20° as a result of Glu428 facing towards the translocation pore cavity in IF state (Fig. S7). The solvation environment shows that Asp316 and Glu419 are connected by three well-ordered water molecules, forming a favorable environment for proton transfer through a Grotthuss mechanism (Fig.4). Furthermore, seven water molecules form a water wire hydrogen network to connect the reoriented Glu428 at the intracellular side and may be possibly involved in the proton transfer from the extracellular to cytoplasmic side of the transporter.

**Figure 4:**
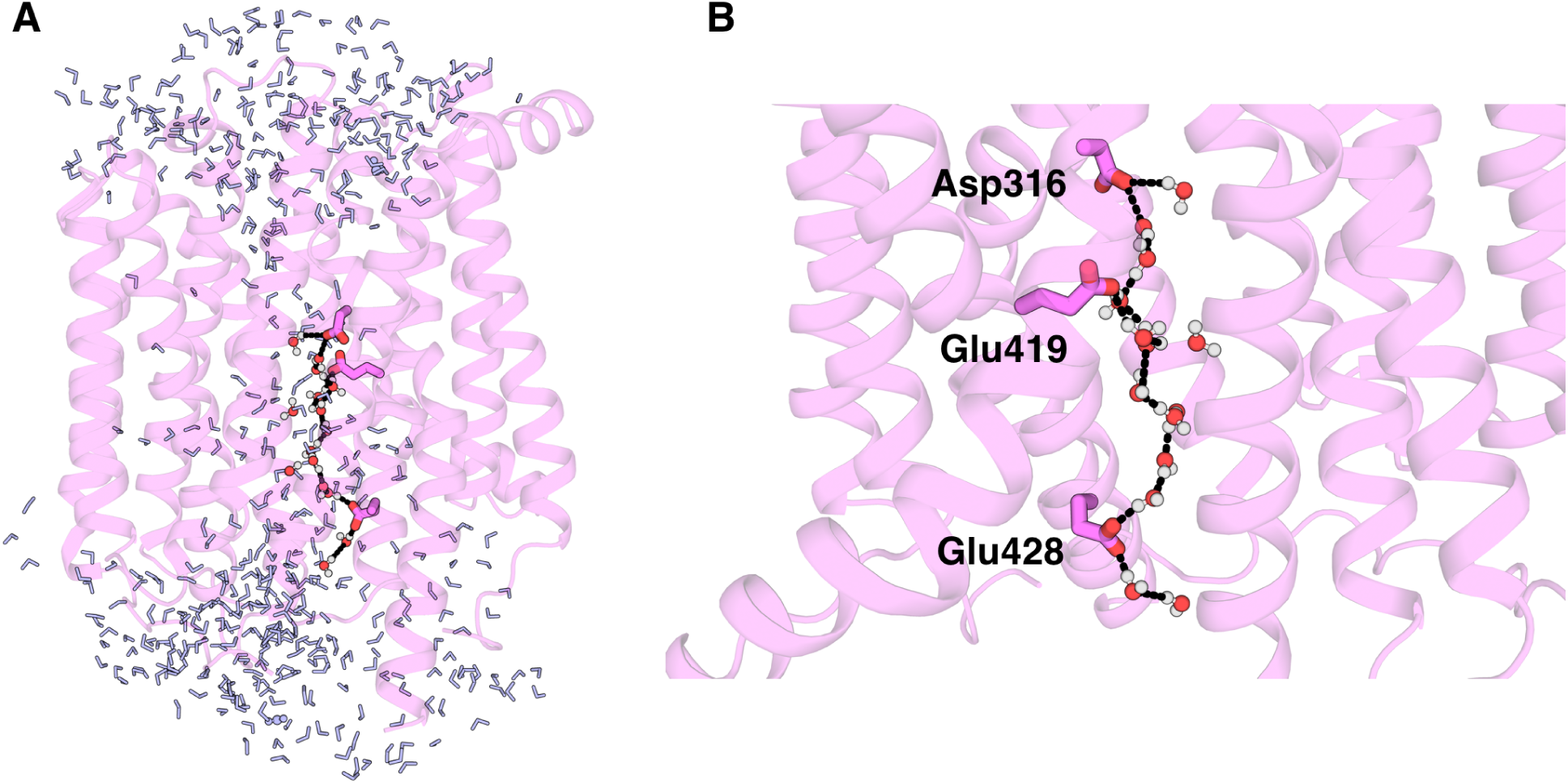
Proton translocation in PepT_So_. A) The water molecules inside the translocation pore of IF state was shown as blue sticks. B) The titratable residues Asp316, Glu419 and Glu428 are shown as stick. The water molecules are represented as ball and stick model. The water molecules form complex network of hydrogen framework, shown as black lines, from extracellular to intracellular side

### Pharmacophore Feature Analysis Guide Drug Design for PepT_So_

Peptide molecules are actively absorbed and transported by intestinal proton-dependent peptide transporter proteins and are considered as valuable tools for improving the pharmacokinetics properties of drug molecules. We extracted the specific pharmacophore features of the translocation pore cavity of intermediate states such OF and IF and mapped the essential features with hPepT_1_ and hPepT_2_ (Fig. S8). The positively charged Arg32 recognizes and binds to a peptide carboxyl group predominately while in the OF state (Fig. 4A). ^61^ The equivalent residue in PepT_2_ is Lys64. Although the residue is different, lysine still carries a positively charged motif and interacts with oppositely charged species (Fig. 5A). Arg25 found in the ExxERxxxY motif acts as a donor and interacts with negatively charged groups in IF state to facilitate peptide transport (Fig. 5B). ^62^ The peptide-bound IF structure of GkPOT shows that phosphate group of Alafosfalin interacts with Arg36 and shows good agreement with our predicted pharmacophore (Fig. S9).^54^ The hydroxyl group of conserved tyrosine (Tyr29, Tyr68, Tyr154, Tyr431) and serine (Ser423) residues, as well as the amide groups of asparagine (Asn157, Asn356) and glutamine (Gln341, Gln147) residues are involved in polar interactions with peptide molecules by acts as an electron donor/acceptors. The scaffolds of drug molecules conjugated with peptide-like molecules form polar contacts with these residues in the translocation pore cavity (Fig. S10, S11). For instance, the prodrug Valacyclovir bound to the IF state shows that carbonyl and ester oxygen atoms exhibit interactions with Tyr41, Tyr79, Tyr163, Asn167 and Asn426 (Fig. S10, S11).^63,64^ The sidechain of Ser161 and Asn454 acts as a hydrogen bond acceptor probes in OF state (Fig. 3C). Our simulation also reveals that peptide backbone (-NH) atoms forms polar contact with the amide group of Asn454. The docked pose of prodrug Valacyclovir in OF state shows that the primary amine group of ligands share polar interactions with Ser161 (Fig. S12). In IF state, Tyr68 and Gln147 act as acceptor moieties.^54^ Although, the glutamine is not conserved, the sidechains of Asn160 and Thr181 have similar chemical properties and may acts as a hydrogen bond acceptor. Another residue, Lys318, carries total positive charge while the equivalent residue is Gln300 and Gln319 in *h*PepT_1_ and *h*PepT_2_, respectively, thereby adopting both donor and acceptor properties. Despite the polar interactions, tyrosine, phenylalanine and tryptophan also play crucial roles in aromatic interaction. Meanwhile aliphatic residues like Ile157 and Leu427 exhibit aliphatic hydrophobic interactions. Our simulation-based, structure optimized pharmacophores may provide a novel insight in designing optimized candidates for improving the bio-oral availability of the peptide like drug molecules.

**Figure 5:**
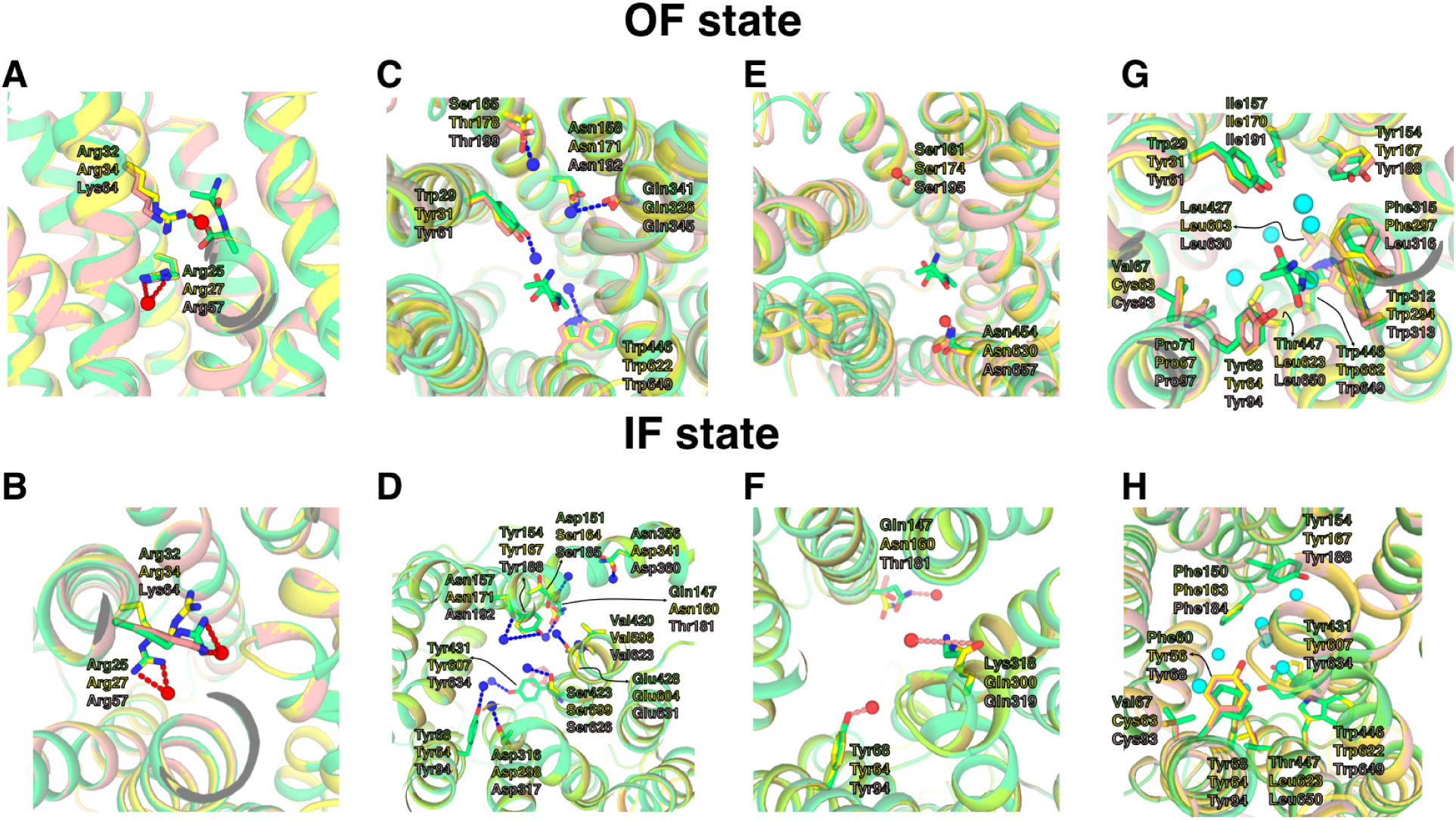
Structure based pharmacophore of A) OF and B) IF state of PepT_So_. The homology models of PepT_1_ and PepT_2_ OF and IF states are mapped to the obtained PepT_So_ pharmacophores. The residues are shown as stick, and pharmacophore features such as a negative ion, hydrogen bond donor, hydrogen bond acceptor, and hydrophobic probes are represented as red, blue, salmon and cyan color spheres, respectively.

## Conclusion

Using extensive simulation, we characterized the Ala-Ala peptide transport by PepT_So_, a human intestinal peptide transporter homologous to PepT_1_. Utilizing Markov State Modeling (MSM), we revealed the free energy landscape, predicting a barrier of approximately 2 kcal/mol for a complete transport cycle. We also identified three kinetically slow states, corresponding to substrate recognition (OF-OC), substrate binding (OC) substrate release (IF) using tIC analysis. These findings sheds light on the energetic requirements and kinetics for peptide translocation across the membrane. Furthermore, we show that substrate binding to the protein modulates its conformational landscape, specifically promoting the transition from the occluded state to the inward-facing state, which is essential for transport. We also used Transition Path Theory (TPT) analysis to decipher the most likely pathways for peptide movement and the critical residues are involved. Specifically, we idenitifed Asn22, Arg32, Glu336 and Gln341 as critical residues for initial recognition, and Gln341, Asp316, His61 and Tyr68 as those for stabilization of the OF state and substrate binding. Finally, by leveraging the structural information obtained from the various intermediate states in our molecular dynamics (MD) simulations, we were able to achieve two key outcomes. Firstly, we identified a potential water network that likely facilitates proton transfer during transport. Secondly, these simulations enabled the determination of pharmacophore features, which can serve as a valuable guide for the rational design of novel drug candidates with enhanced efficacy. Future investigations could delve deeper by exploring the translocation mechanisms of more complex and therapeutically relevant peptides, potentially including those with varying lengths, charges, and functionalities. Additionally, investigating the effects of mutations on the conformational dynamics of PepT_So_ with different substrates would provide valuable insights into potential disease mechanisms associated with dysfunctional peptide transport. Overall, our findings offer novel insights into the interplay between conformational dynamics, substrate transport, and proton transport in PepT_So_, which may helps in rational drug design for improving drug efficacy.

## Acknowledgement

D.S. acknowledges support from NIGMS MIRA award R35GM-142745 and NSF Early CAREER Award (MCB-1845606). We thank the Blue Waters sustained-petascale computing project, which is supported by the National Science Foundation and the state of Illinois.

## Data Availability

Molecular dynamics simulations trajectories have been uploaded to https://uofi.box.com/s/8f9sh1o0rdw6xiwo76cls0bsokkl2egg.

