## Supplementary Information for "Energetics of substrate transport in proton-dependent oligopeptide transporters"

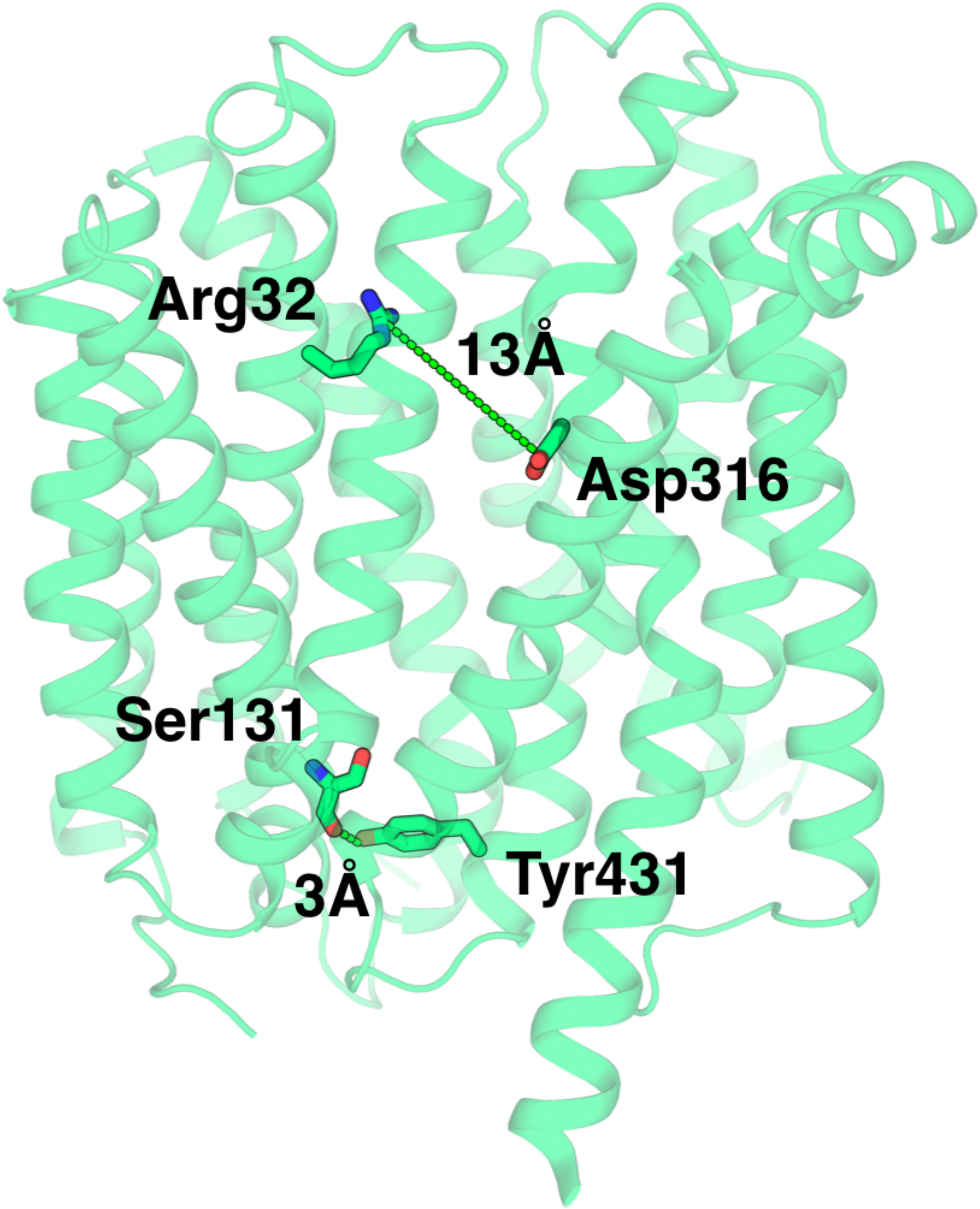


Figure S1: The distance between the extracellular and intracellular gating residues. The residue pair distance is used for the projection of the conformation landscape plots.

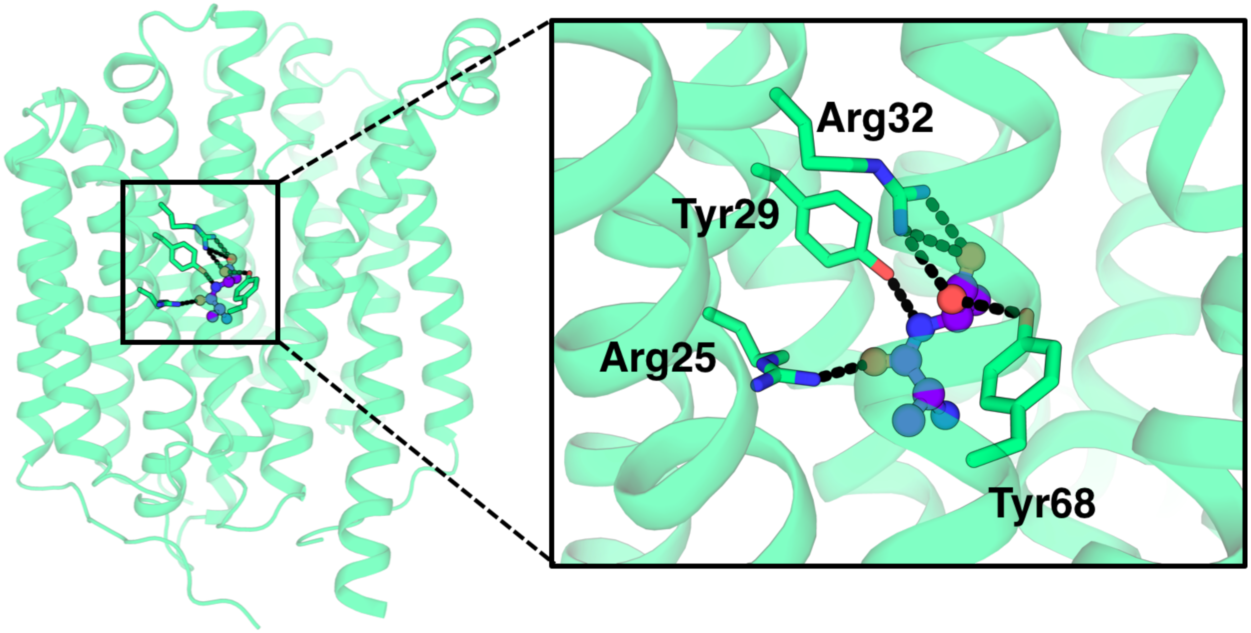


Figure S2: Ala-Ala peptide form network of polar contacts in the translocation pore channel in the OF state. Tyr29 and Tyr68 residues play crucial role in diffusion of the substrate molecules to inside the intracellular cavity and the conformational changes.

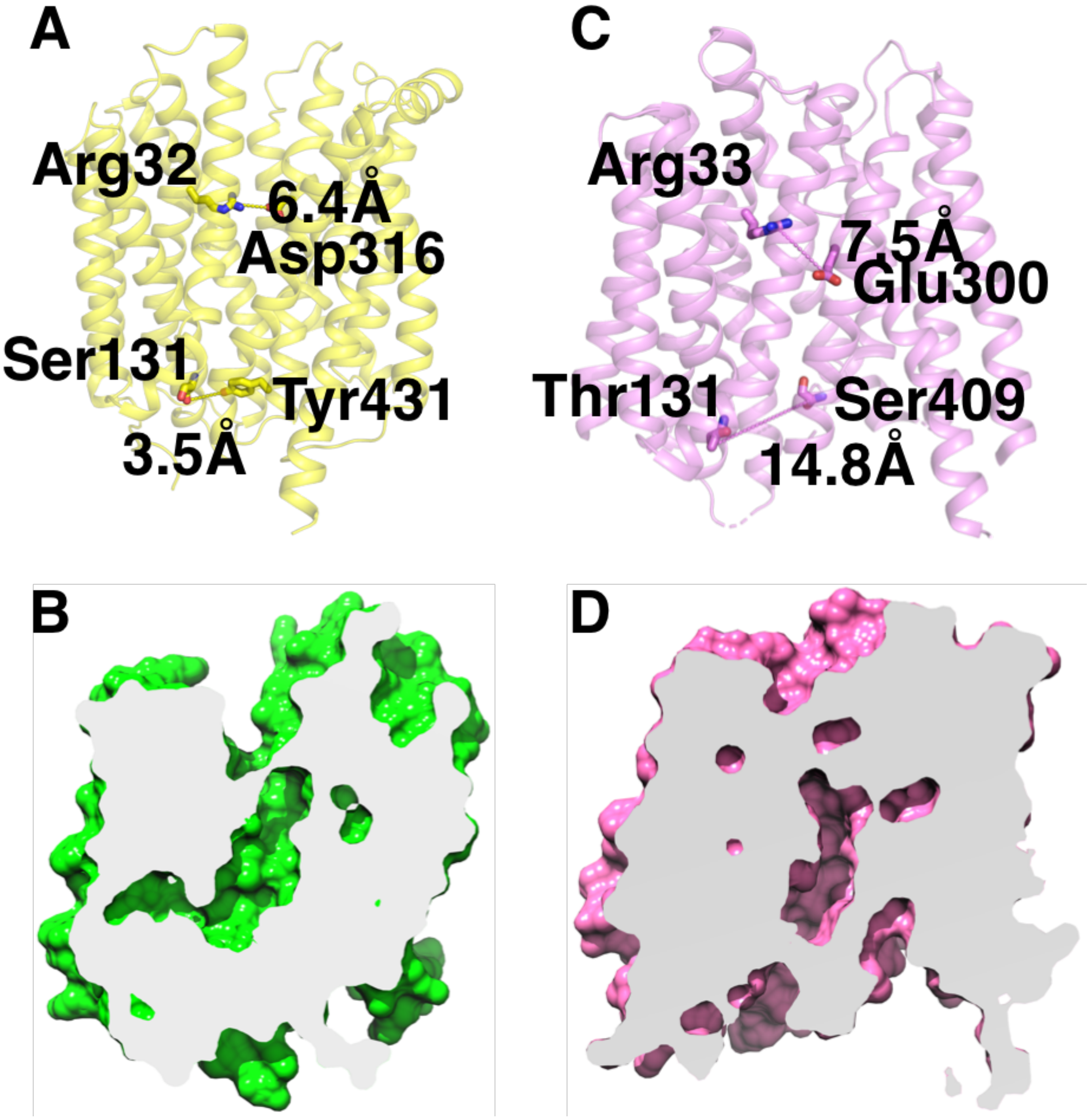


Figure S3: The comparison of predicted OC state of $PepT_{So}$ and $PepT_{St}$ . A) and C) The OC states of $PepT_{So}$ was shown as carton and surface, respectively. B) and D) The OC states of $PepT_{St}$ was shown as carton and surface, respectively

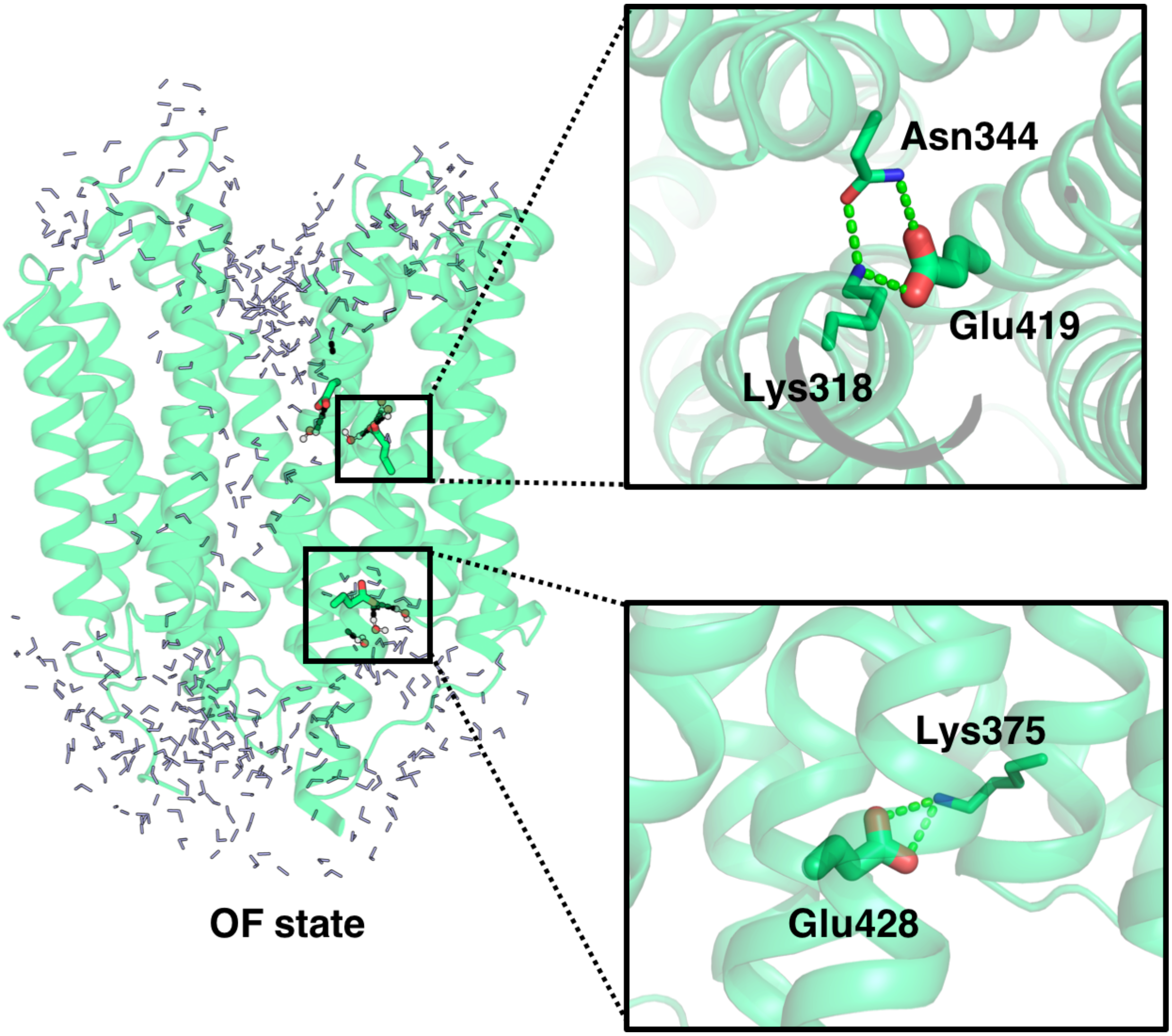


Figure S4: The water filled pore cavity of the OF state. Glu419 was buried between the helices 10 and 8. The polar residues Asn344 and Lys318 interacts with charged sidechain of Glu419. Another titratable residue, Glu428 interacts with Lys375 at the intracellular side

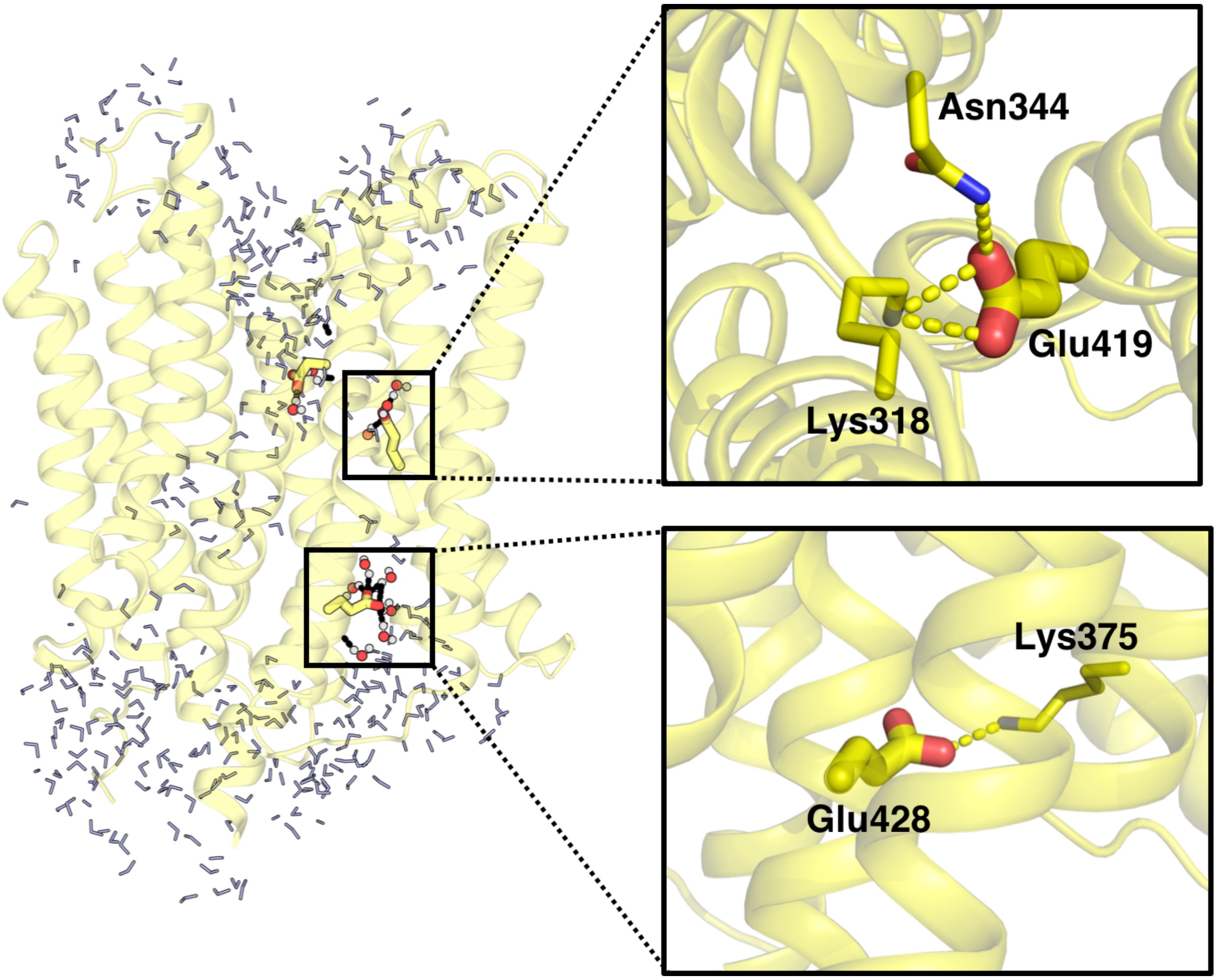


Figure S5: The water filled pore cavity of the OC state. Glu419 was buried between the helices 10 and 8. The polar residues Asn344 and Lys318 interacts with charged sidechain of Glu419. Another titratable residue, Glu428 interacts with Lys375 at the intracellular side.

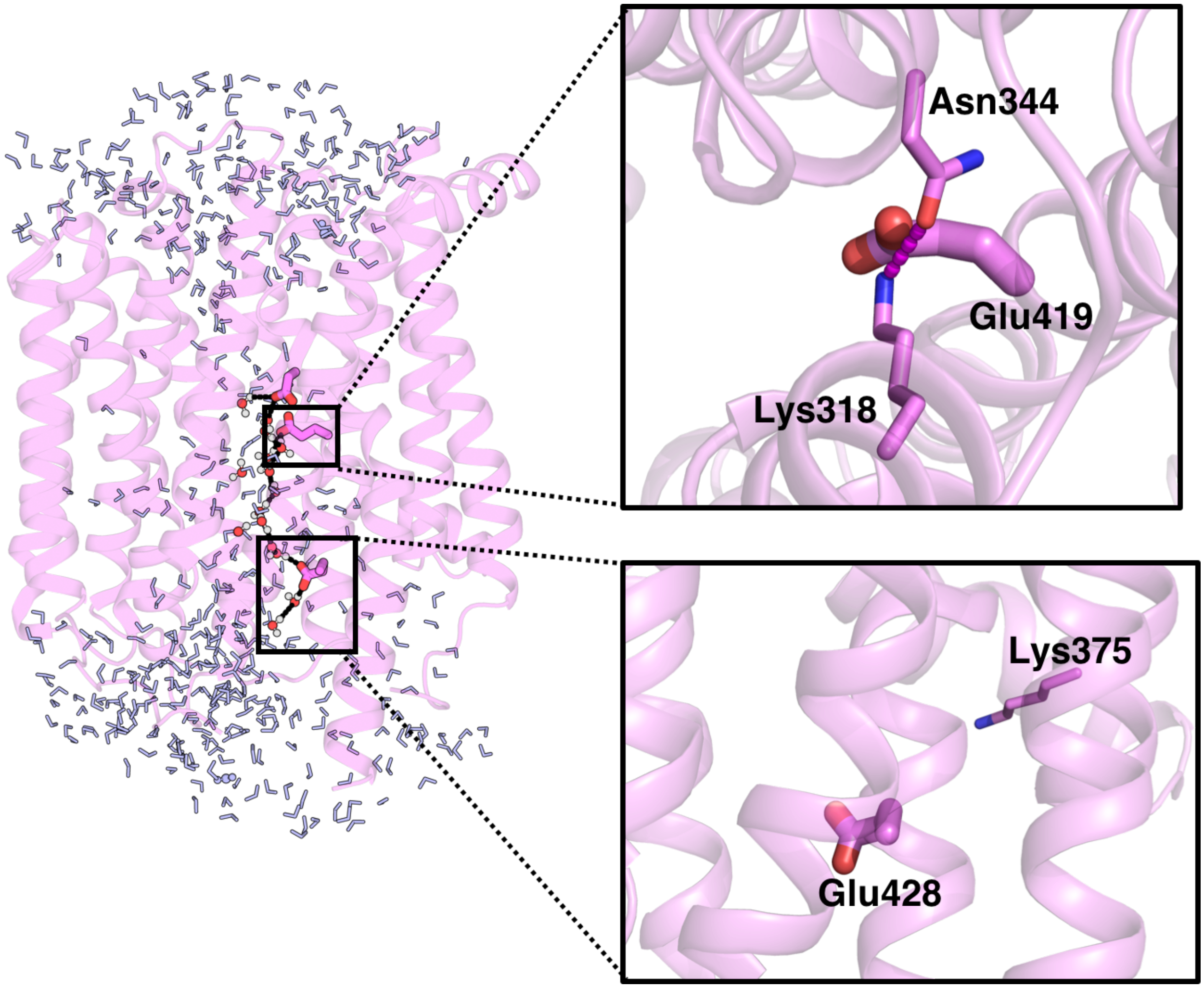


Figure S6: The water filled pore cavity of the IF state. Glu419 was exposed to the transport pore channel and the kinked helices leads to the formation of Lys318 and Asn344 interaction. The titratable residue, Glu428 also faces the pore cavity at the intracellular side as the helix rotates up to 20° during the conformational transition to IF.

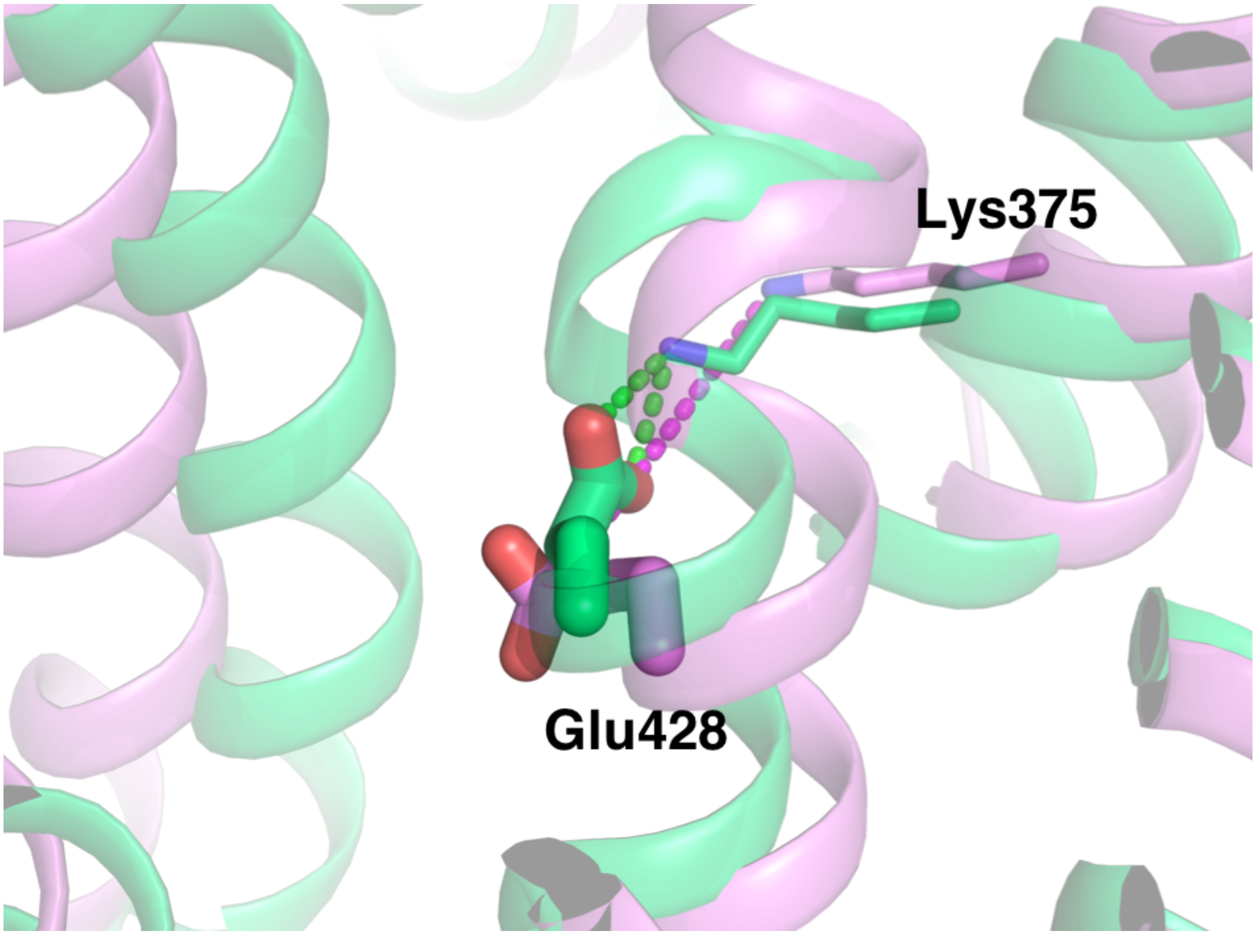


Figure S7: The differences in sidechain orientation of Glu428 in OF and IF state. In IF state, Lys375 move outward thereby loses interaction with Glu428.

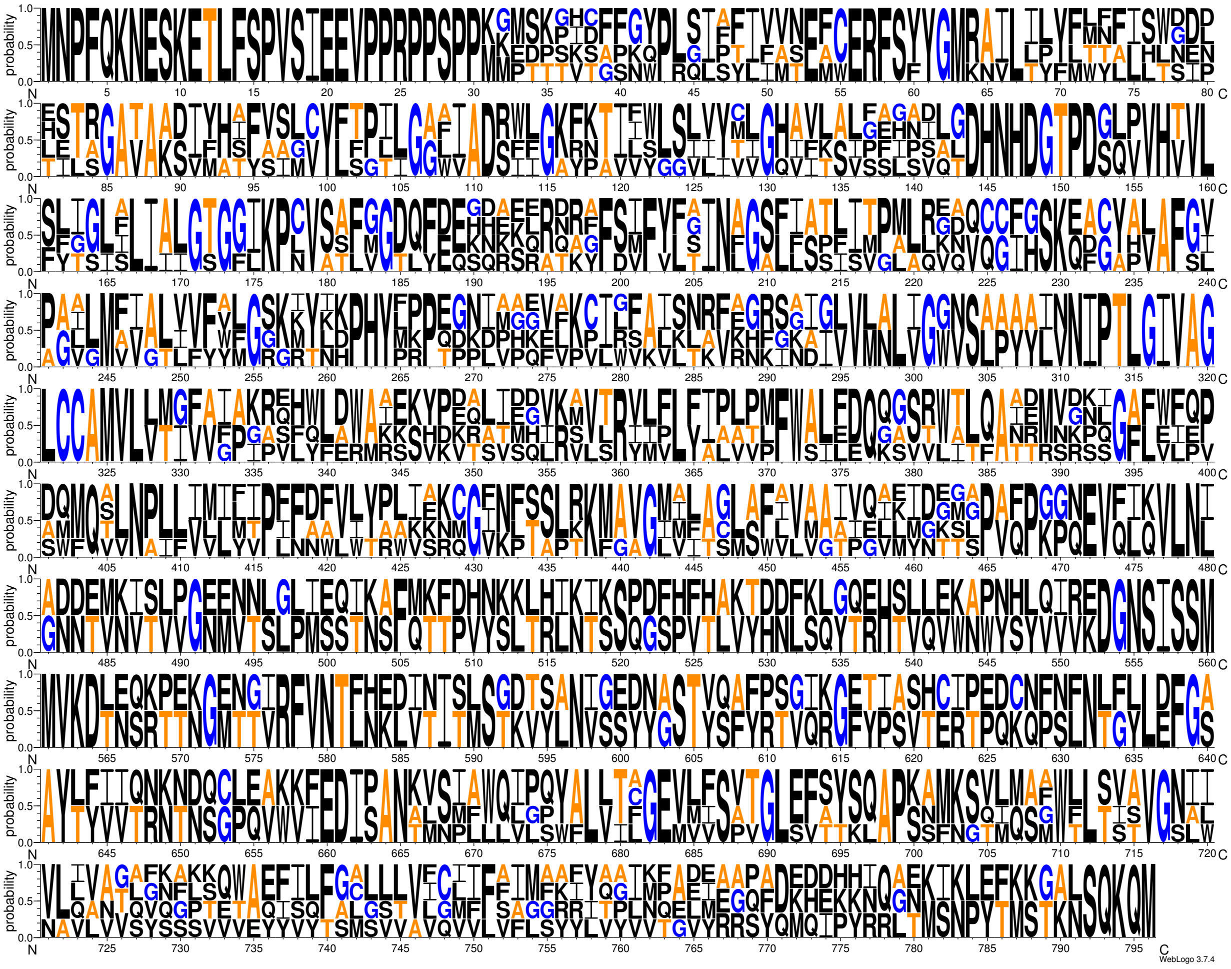

Figure S8: Multiple sequence alignment of $PepT_{So}$, $PepT_1$ and $PepT_2$. The sequence alignment was shown as WebLogo format.

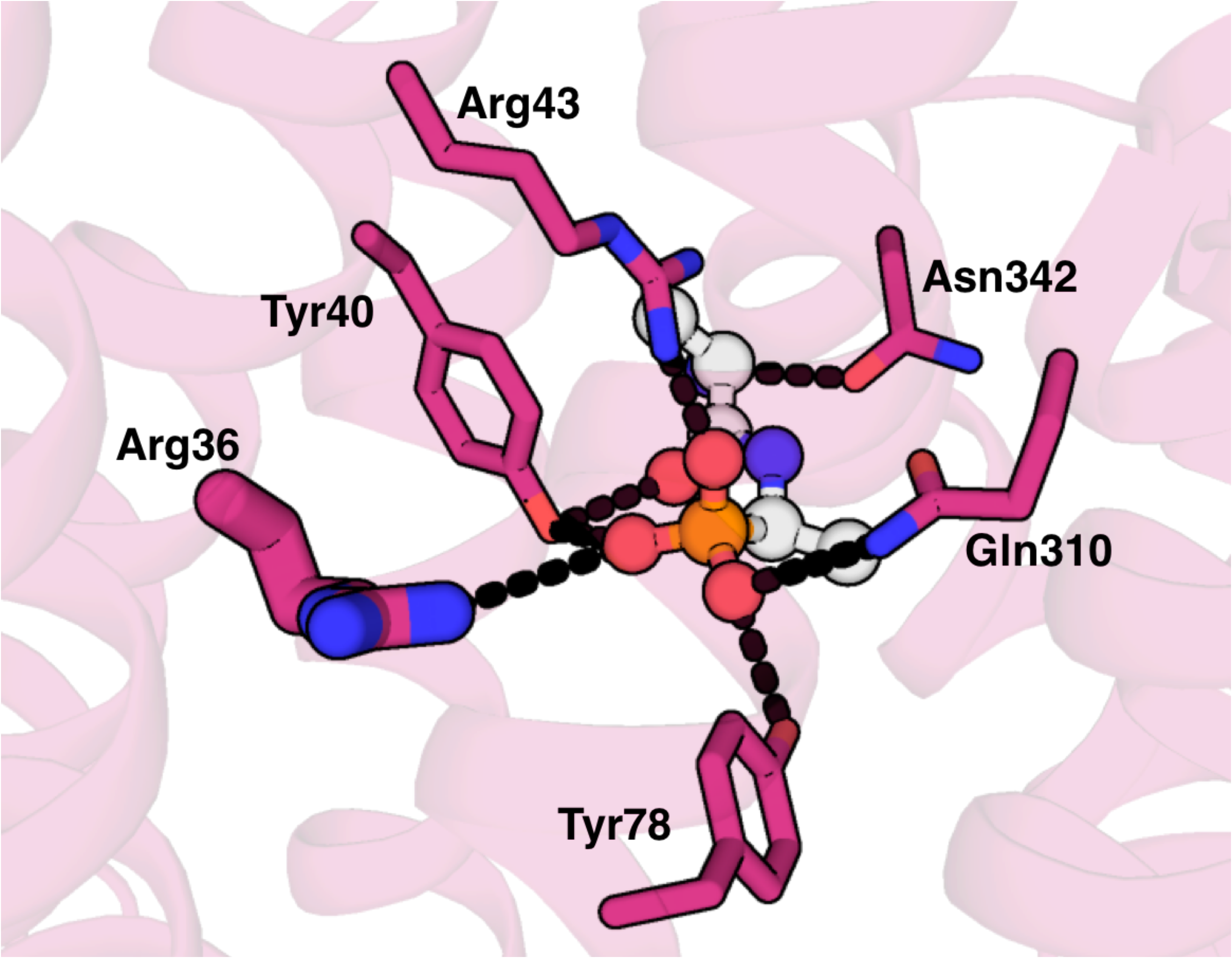


Figure S9: Alafosfalin bound IF state crystal structure of GkPOT.[1] Alafosfalin was shown as ball and stick model and the residues that form polar interactions are shown in sticks, respectively.

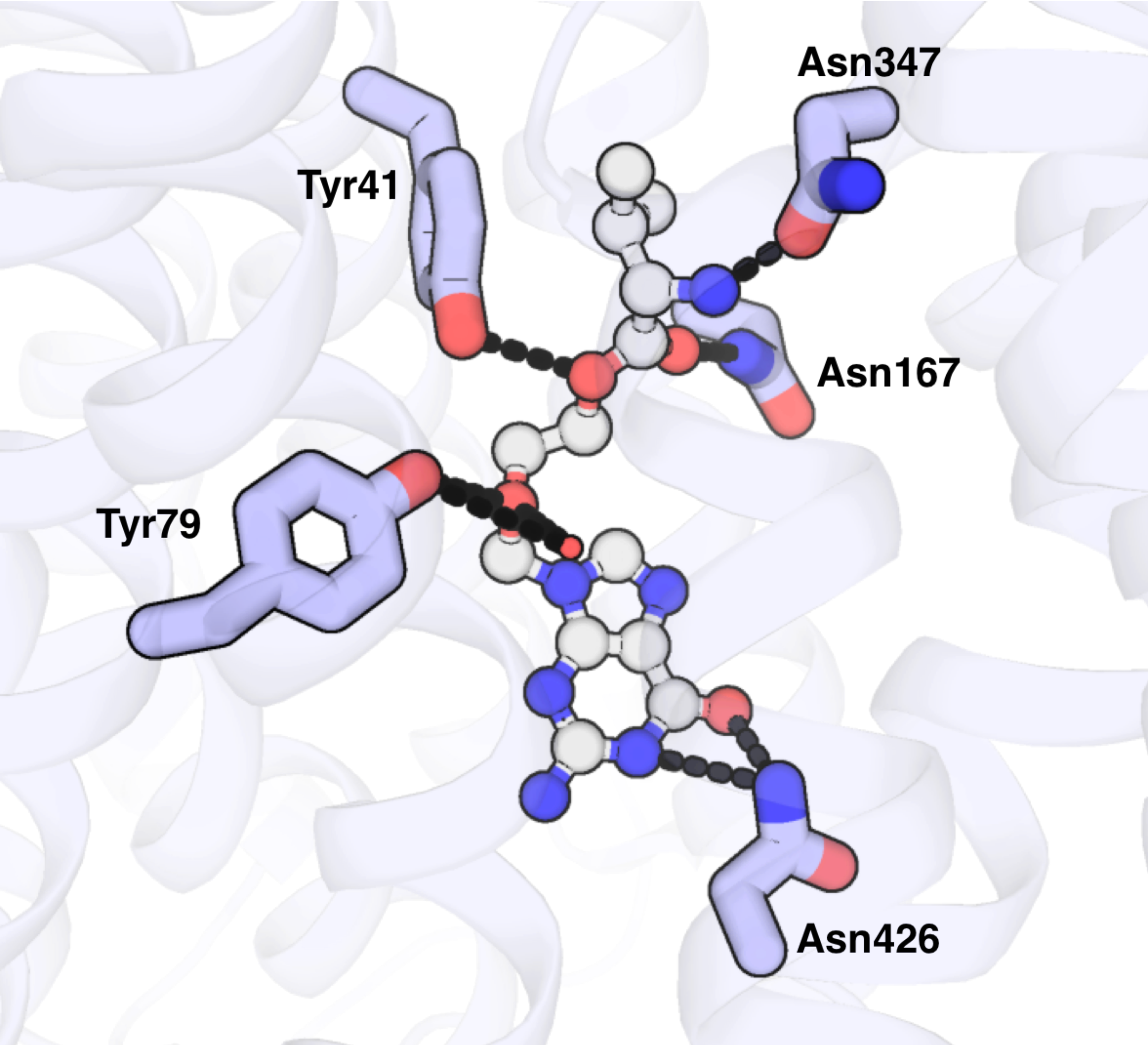


Figure S10: Prodrug, Valacyclovir bound IF state crystal structure of $PepT_{Sh}$.[2] Valacyclovir was shown as ball and stick model and the residues that form polar interactions are shown in sticks, respectively.

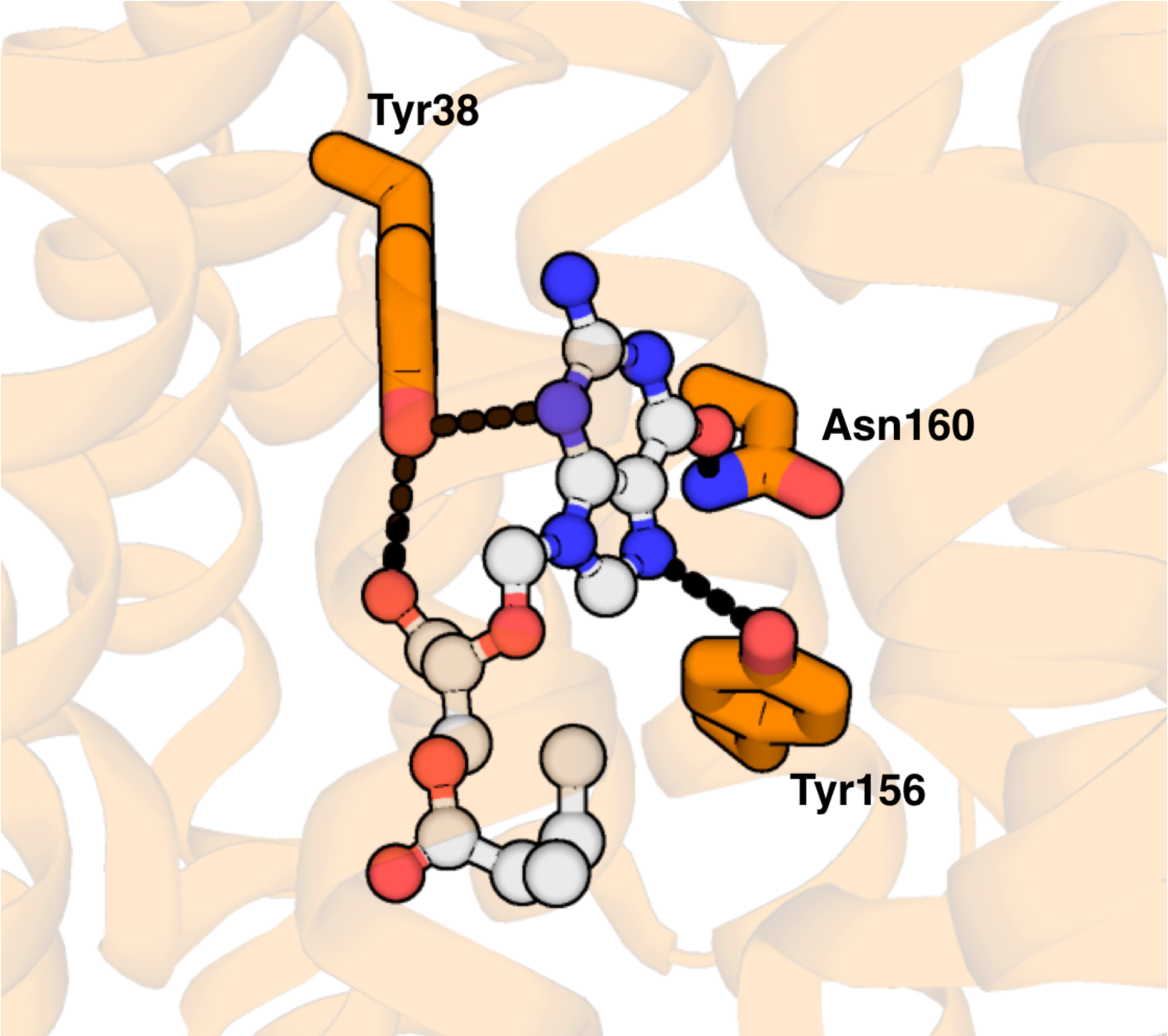


Figure S11: Prodrug, Valacyclovir bound IF state crystal structure of DtpA from E.coli.[3] Valacyclovir was shown as ball and stick model and the residues that form polar interactions are shown in sticks, respectively.

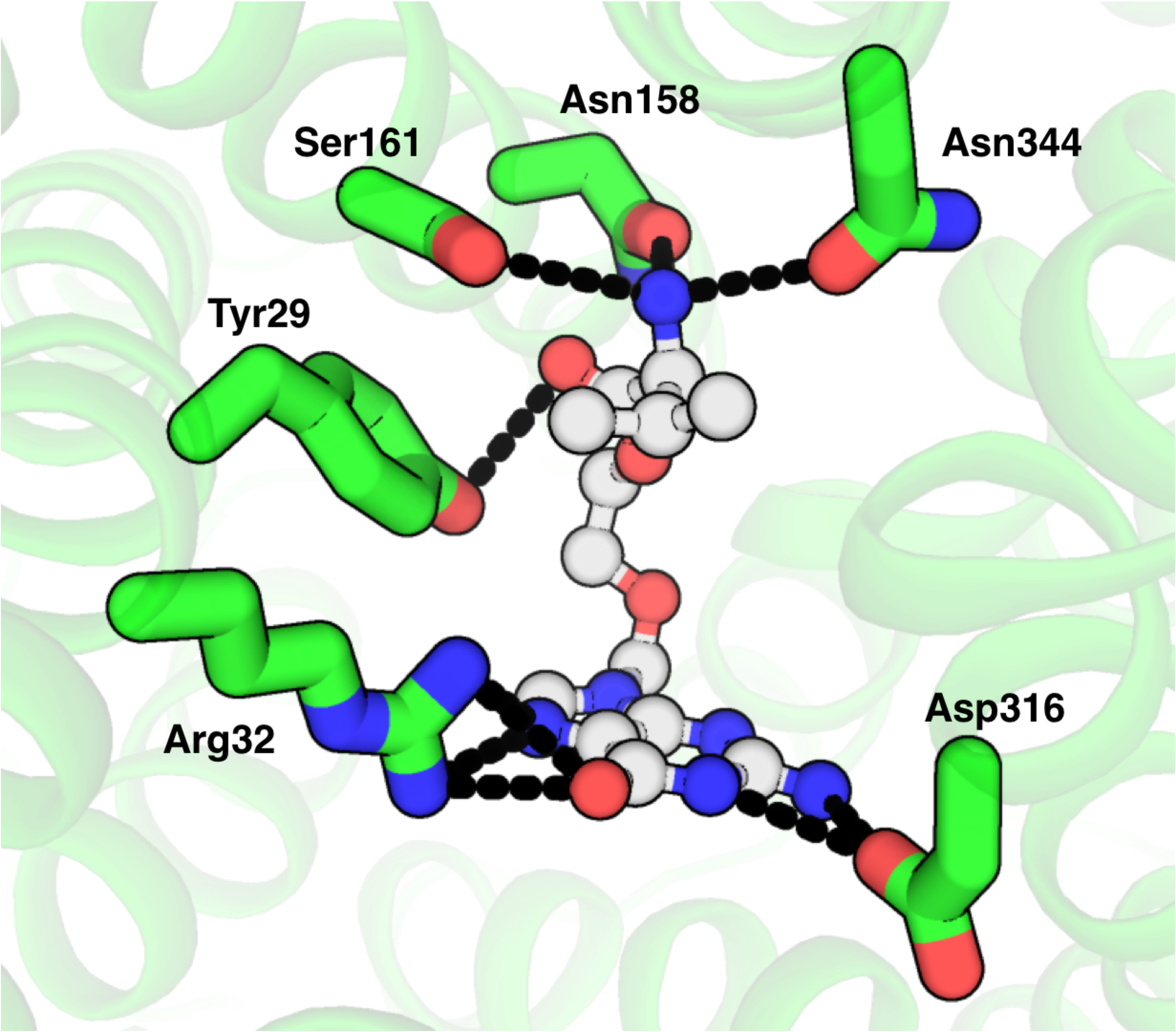


Figure S12: Prodrug, Valacyclovir docked to the OF state of $PepT_{So}$. Valacyclovir was shown as ball and stick model and the residues that form polar interactions are shown in sticks, respectively.

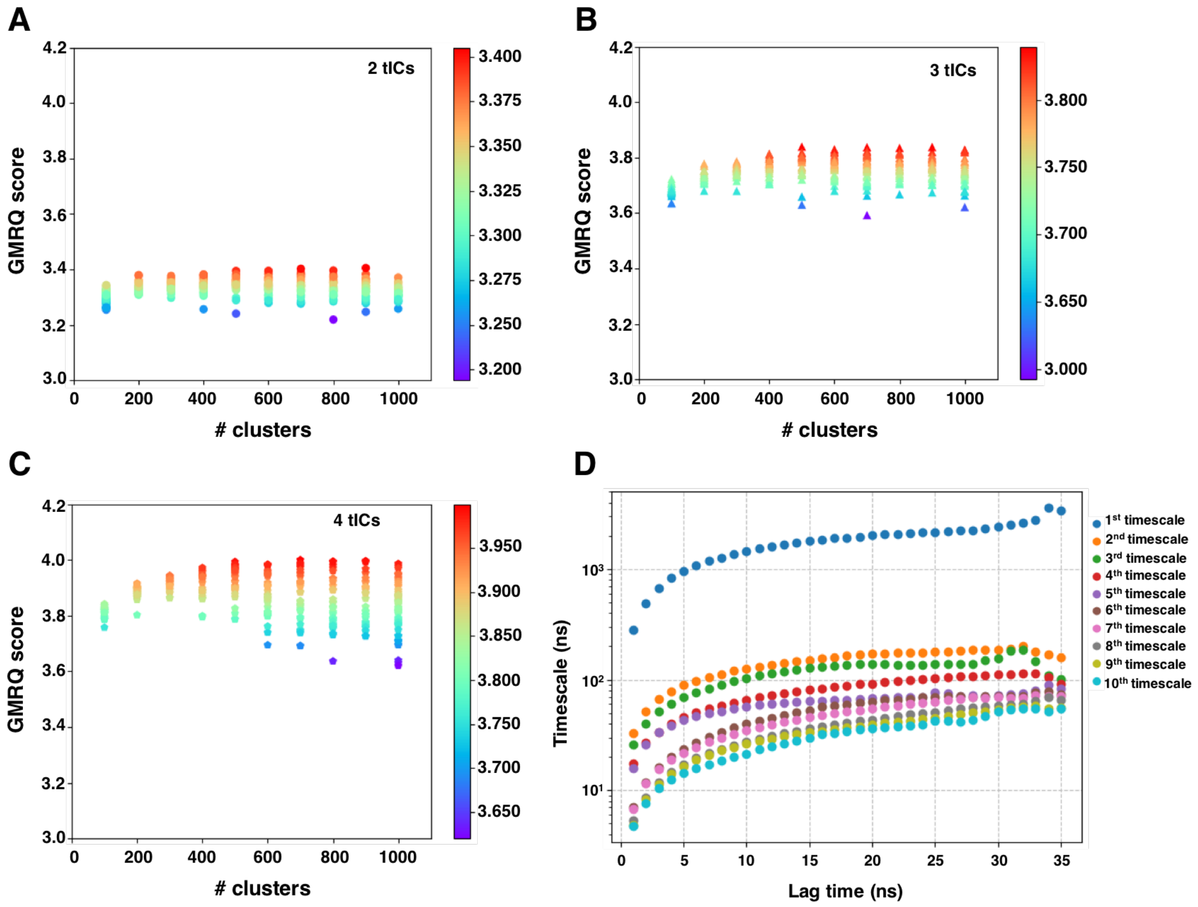


Figure S13: Osprey analysis and Implied timescale plot of PepT$_{So}$.[4] The osprey plots for A) 2tICs, B) 3tICs and C) 4tICs are shown, respectively. The high scoring 4tICs and 500 clusters are chosen for constructing the MSM. The lag time of 20 ns was used for the MSM.

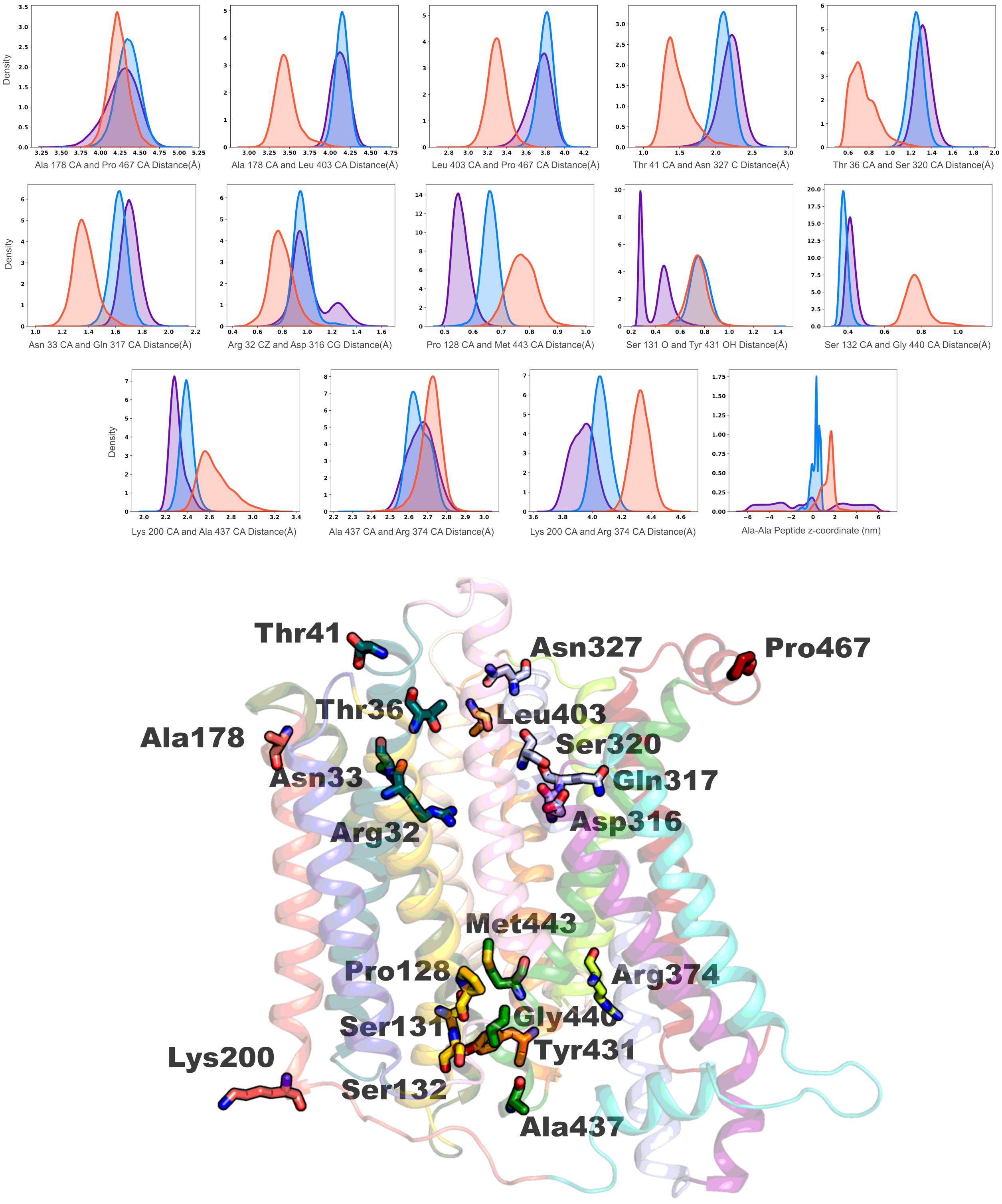


Figure S14: Features used to construct time-independent components (tICs). The three colors corresponds to the color of the minima of the three kinetically slow states on the tIC landscape. The structure of $PepT_{So}$ with the residues involved in all features shown in sticks and labeled.

Table S1: Adaptive sampling rounds for simulation.

| Round | Time ($\mu$s) |
|---|---|
| 1 | 0.3 |
| 2 | 0.9 |
| 3 | 0.9 |
| 4 | 0.7 |
| 5 | 1.0 |
| 6 | 1.0 |
| 7 | 0.7 |
| 8 | 0.9 |
| 9 | 0.4 |
| 10 | 1.0 |
| 11 | 0.5 |
| 12 | 0.7 |
| 13 | 2.4 |
| 14 | 4.7 |
| 15 | 4.8 |
| 16 | 2.4 |
| 17 | 2.5 |
| 18 | 1.3 |
| 19 | 1.0 |
| 20 | 5.1 |
| 21 | 3.1 |
| 22 | 3.1 |
| 23 | 3.6 |
| 24 | 3.7 |
| 25 | 3.6 |
| 26 | 1.0 |
| 27 | 2.1 |
| 28 | 1.5 |
| 29 | 1.6 |
| 30 | 10.3 |
| 31 | 3.6 |
| 32 | 1.9 |
| 33 | 3.0 |
| 34 | 9.8 |
| **Total** | 85.2 |
